# ZDHHC14-Mediated Palmitoylation of TBK1 Promotes Pathological Cardiac Hypertrophy via Type I Interferon Signaling Activation

**DOI:** 10.64898/2026.07.24.740659

**Authors:** Wei Zhao, Ying Yang, Gaoyuan Ge, Ruochen Xu

## Abstract

Pathological cardiac hypertrophy represents a common maladaptive response to cardiovascular stress and constitutes a major harbinger of heart failure. Although S-palmitoylation—a reversible post-translational modification—critically governs protein localization, trafficking, and stability, its involvement in cardiac hypertrophy remains poorly characterized. In this study, we aimed to explore the role and regulatory mechanism of a palmitoyltransferase, zinc finger DHHC-type palmitoyltransferase 14 (ZDHHC14), in cardiac hypertrophy. We found that ZDHHC14 was significantly upregulated in cardiac hypertrophy tissues from both human patients and mouse models. Cardiomyocyte-specific ZDHHC14 knockdown ameliorated transverse aortic constriction (TAC)-induced cardiac hypertrophy and dysfunction in male mice, whereas cardiac-specific ZDHHC14 overexpression via AAV9 exacerbated these pathological phenotypes. Mechanistically, TANK-binding kinase 1 (TBK1) was identified as a novel substrate of ZDHHC14 through interactomic screening. ZDHHC14 catalyzed TBK1 palmitoylation at cysteine 267, which in turn facilitated TBK1 phosphorylation and subsequent activation of type I interferon (IFN-I) signaling, ultimately promoting cardiac hypertrophy. Importantly, our findings demonstrate that the TBK1-C267S mutation rectifies ZDHHC14 overexpression-induced exacerbation of cardiac hypertrophy. This study illustrated a ZDHHC14-TBK1-IFN-I axis in regulating cardiac hypertrophy, which may provide a potential therapeutic target to ameliorate pathological cardiac hypertrophy.

## Introduction

Pathological cardiac hypertrophy represents a maladaptive response commonly arising from chronic hypertension and constitutes a critical precursor to heart failure (HF), the end-stage manifestation of cardiovascular disease^1^. HF remains a leading cause of mortality and morbidity worldwide, imposing a substantial burden on public health^2, 3^. Thus, halting or retarding the progression of pathological hypertrophy offers a viable avenue for averting the eventual transition to overt HF^4^. Although contemporary therapeutic modalities—including pharmacotherapy, surgical procedures, device-based interventions, and lifestyle modifications—are widely available, only a limited subset of patients achieves meaningful reversal of hypertrophic remodeling^5^. This clinical reality underscores the urgent demand for the identification of precise molecular targets that can inform the development of novel and more effective treatment paradigms.

A diverse array of intracellular signaling cascades has been implicated in the pathogenesis of pathological cardiac remodeling and the subsequent development of heart failure^6, 7^. Among these pathways, type I interferon (IFN-I) signaling has emerged as a critical contributor to maladaptive cardiac hypertrophy^8, 9^. Tank-binding kinase 1 (TBK1) serves as a pivotal effector within this signaling axis, functioning as an essential kinase that catalyzes interferon regulatory factor 3 (IRF3) phosphorylation upon recruitment by upstream adaptors—including the interferon gene stimulator (STING), mitochondrial antiviral signaling protein (MAVS), and the TIR-domain-containing adapter-inducing interferon-β (TRIF)-dependent cascade—thereby culminating in the induction of type I interferons^10, 11^. Of particular interest, accumulating evidence highlights the importance of TBK1 and its phosphorylation status in the context of pathological cardiac remodeling^12^. Moreover, the activation state and biological functions of TBK1 are subject to intricate regulation via multiple post-translational modifications (PTMs), encompassing ubiquitination, phosphorylation, hydroxylation, and acetylation^13–15^. Accordingly, pharmacological or genetic modulation of these TBK1 PTMs represents a viable therapeutic avenue for either potentiating or attenuating TBK1-mediated IFN-I signaling. There are hundreds of protein PTMs identified to date^16^. Identifying the critical PTMs of TBK1 and its functional role in pathological cardiac hypertrophy represents an urgent problem to be solved.

PTMs serve as fundamental mechanisms governing protein functionality and subcellular distribution. Among these, S-palmitoylation represents a reversible lipidation process mediated by a family of 23 S-acyltransferases, all of which share a conserved zinc-finger aspartate-histidine-histidine-cysteine (zDHHC) motif^17^. The zDHHC S-acyltransferases are integral membrane proteins possessing multiple transmembrane domains; although they predominantly reside in the endoplasmic reticulum and Golgi apparatus, certain members are also found at the plasma membrane, throughout the endomembrane system, or associated with intracellular vesicles^18^. Although substrate overlap exists among certain zDHHC isoforms, these enzymes generally exhibit pronounced selectivity and stringent substrate recognition, largely dictated by recruitment domains situated within their cytoplasmic regions^19^. Protein S-palmitoylation exerts diverse functional effects, including enhanced protein stability, redirected intracellular localization, and altered biological activity^20^. Palmitoylated proteins have been implicated in the pathogenesis and progression of numerous human disorders, spanning malignancies^21^, immune-inflammatory responses^22^, metabolic diseases^23^, and colitis^24^. Nevertheless, current understanding of zDHHC enzymes and S-palmitoylation within the cardiovascular system remains predominantly confined to their roles in ion channel modulation and electrophysiological properties^19, 25^. Notably, their involvement in cardiomyocyte signaling cascades, hypertrophic remodeling, and the development of heart failure has received comparatively little attention.

Zinc finger DHHC-type palmitoyltransferase 14 (ZDHHC14), a key member of the palmitoyl acyltransferase (PAT) family, catalyzes the S-palmitoylation of various protein substrates, modulating a wide array of cellular processes—ranging from ferroptosis^26^ and cell cycle regulation^27^ to metabolic reprogramming^28^ and ion channel homeostasis^29^. Through these multifaceted actions, ZDHHC14 has been implicated in the pathogenesis of both malignancies and neurological disorders. However, the role and underlying mechanism of ZDHHC14 in pathological cardiac hypertrophy have not yet been reported.

Integrated transcriptomic profiling coupled with functional validation revealed a marked upregulation of the palmitoyl acyltransferase ZDHHC14 in heart tissues obtained from both hypertrophic murine models and human patients with cardiac hypertrophy. Through cardiomyocyte-restricted manipulation of ZDHHC14 expression—via targeted knockdown and forced overexpression—we demonstrated that ZDHHC14 knockdown effectively attenuated TAC-induced myocardial hypertrophy and cardiac dysfunction, whereas its overexpression significantly aggravated these pathological manifestations. Mechanistically, ZDHHC14 interacts with and mediates TBK1 palmitoylation, thereby promoting TBK1 phosphorylation and the activation of the IFN-I signaling. We also found that ZDHHC14 palmitoylates TBK1 at C267 site, and TBK1-C267S mutant ameliorated ZDHHC14 overexpression-induced exacerbation of cardiac hypertrophy. Therefore, inhibiting the ZDHHC14-TBK1 axis might represent a promising therapeutic strategy for treatment of cardiac hypertrophy.

## RESULTS

### ZDHHC14 is significantly increased in both mice and humans exhibiting cardiac hypertrophy

Recent studies has demonstrated that palmitoyl acyltransferases (PATs) exert critical regulatory roles across a panel of cardiovascular diseases^30^. However, the contribution and underlying molecular mechanism of PATs in pathological cardiac hypertrophy remain poorly characterized. By analyzing the differentially expressed genes (DEGs) in myocardial tissues from patients with hypertrophic cardiomyopathy (HCM) and healthy controls within the GSE141910 dataset, we intersected the DEGs with the PATs, and identified that the expression levels of ZDHHC9, ZDHHC11, ZDHHC14 and ZDHHC23 were significantly altered (Fig.1A-B). Subsequent validation experiments in established mouse models of cardiac hypertrophy further confirmed that only ZDHHC14 exhibited a remarkable upregulation in the myocardial tissues of hypertrophic mice (Fig.1C). Further, we confirmed that ZDHHC14 is significantly up-regulated in multiple human cardiac hypertrophy datasets (Fig.1D-H and Fig.S1A). Similarly, the expression of ZDHHC14 also rises in the myocardial tissues of patients with dilated cardiomyopathy (DCM) and ischemic cardiomyopathy (ICM) recorded in public GEO databases (Fig.S1A-D). Subsequent experiments verified that both the mRNA and protein levels of ZDHHC14 are upregulated in human hypertrophic heart tissues, compared with the normal control groups (Fig.1I-J, Fig.S1E). Furthermore, the mRNA and protein expression of ZDHHC14 was also increased in heart tissues from both Ang II- and TAC- induced cardiac hypertrophy models (Fig.1K-L, Fig.S1F).

**Fig. 1.**
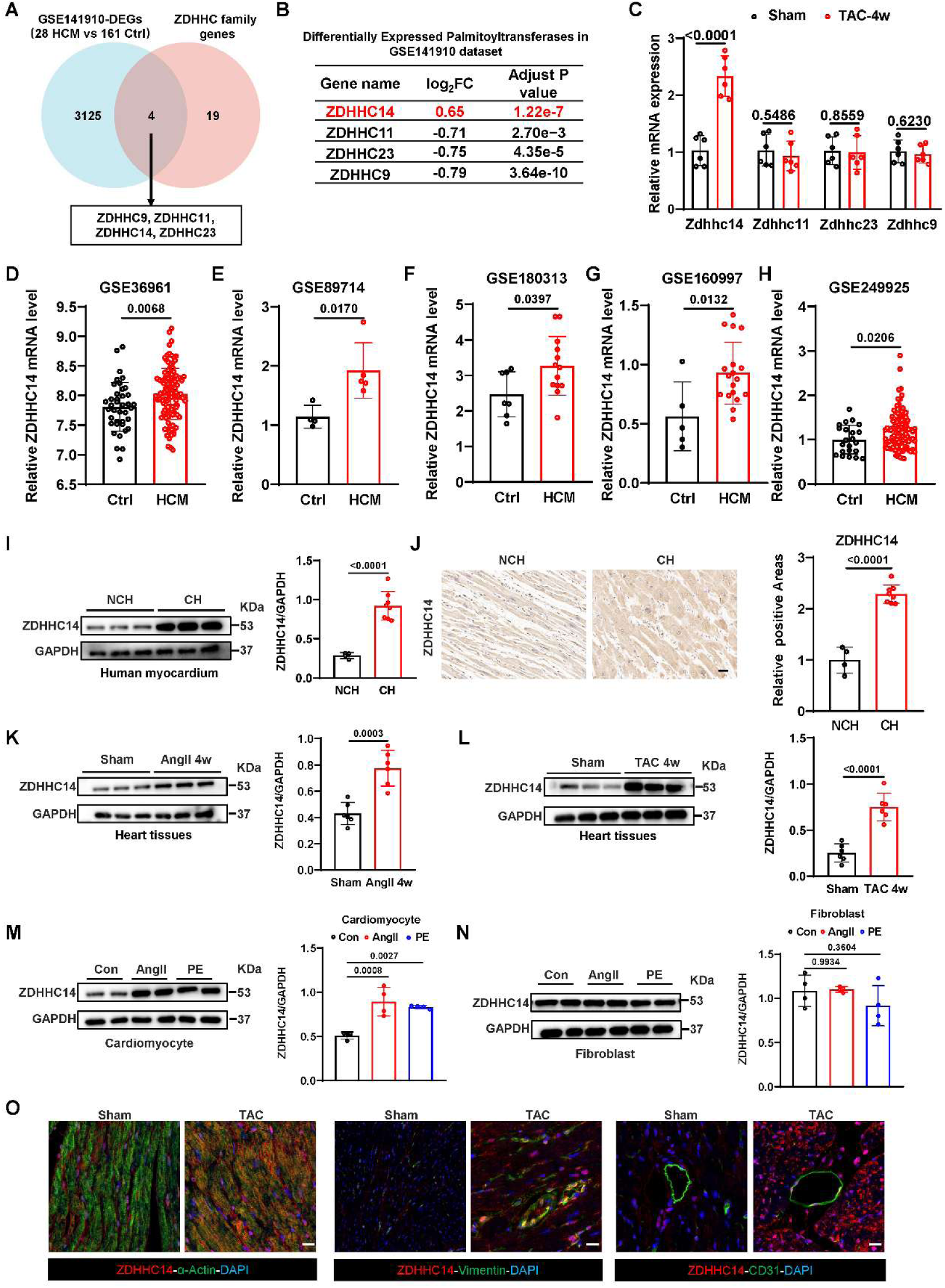
Identification of ZDHHC14 as an upregulated factor in mouse and human cardiac hypertrophy. A. Integrating the differently expressed genes (DEGs) from GEO database (GEO accession no. GSE141910) of human cardiac hypertrophy (n = 161 biological replicates for ctrl and n = 28 biological replicates for hypertrophic cardiomyopathy (HCM) patients) and known 23 palmitoyl acyltransferases. B. The differently expressed palmitoyl acyltransferases between control patients (n = 161 biological replicates) and HCM patients (n = 28 biological replicates) from GEO database (GEO accession no. GSE141910). C. RT-qPCR analysis of ZDHHC9, ZDHHC11, ZDHHC14, and ZDHHC23 mRNA level in heart tissues from sham (n = 6 biological replicates) and TAC mice (n = 6 biological replicates). D. The mRNA expression of ZDHHC14 in heart tissues from control patients (n = 39 biological replicates) and HCM patients (n = 106 biological replicates) in the public RNA-Sequencing database (GEO accession no. GSE36961). E. The mRNA expression of ZDHHC14 in heart tissues from control patients (n = 4 biological replicates) and HCM patients (n = 5 biological replicates) in the public RNA-Sequencing database (GEO accession no. GSE89714). F. The mRNA expression of ZDHHC14 in heart tissues from control patients (n = 7 biological replicates) and HCM patients (n = 13 biological replicates) in the public RNA-Sequencing database (GEO accession no. GSE180313). G. The mRNA expression of ZDHHC14 in heart tissues from control patients (n = 5 biological replicates) and HCM patients (n = 18 biological replicates) in the public RNA-Sequencing database (GEO accession no. GSE160997). H. The mRNA expression of ZDHHC14 in heart tissues from control patients (n = 22 biological replicates) and HCM patients (n = 97 biological replicates) in the public RNA-Sequencing database (GEO accession no. GSE249925). I. Representative immunoblot of ZDHHC14 in human hypertrophic myocardium (CH, cardiac hypertrophy; NCH, no cardiac hypertrophy; n = 4 biological replicates for NCH and n = 8 biological replicates for CH). J. Representative images and quantitation of immunohistochemical staining of ZDHHC14 in human heart tissues from NCH (n = 4 biological replicates) and CH patients (n = 8 biological replicates). Scale bar, 20μm. K. ZDHHC14 protein level measured by western blotting in heart tissues from sham mice (n = 6 biological replicates) and Ang II-induced cardiac hypertrophy mice (n = 6 biological replicates). L. ZDHHC14 protein levels measured by western blotting in heart tissues from sham mice (n = 6 biological replicates) and TAC-induced cardiac hypertrophy mice (n = 6 biological replicates). M. ZDHHC14 protein levels measured by western blotting in primary cardiomyocytes stimulated with Ang II (1 μM), PE(50 μM) or PBS for 24 h (n = 4 biological replicates). N. ZDHHC14 protein levels measured by western blotting in primary fibroblasts stimulated with Ang II (1 μM), PE(50 μM) or PBS for 24 h (n = 4 biological replicates). O. The cellular origin of ZDHHC14 in heart sections in sham mice (n = 6 biological replicates) and TAC mice (n = 6 biological replicates) were assessed using immunofluorescence staining. Scale bar, 20 μm. Red: ZDHHC14; Green: actinin for cardiomyocyte; vimentin for fibroblast; CD31 for endothelial cell. For statistical analysis, two-tailed unpaired student t-test was used for (C-L), and one-way ANOVA/Tukey test was used for (M and O). Data are presented as mean ± SEM. P < 0.05 indicates a statistically significant difference.

In accordance with our previously observations, the mRNA and protein levels of ZDHHC14 were significantly upregulated in neonatal rat primary cardiomyocytes (NRCMs) following stimulation with phenylephrine (PE) or Ang II, as illustrated in Fig.1M and Fig.S1G. Notably, no comparable induction of ZDHHC14 expression was detected in isolated primary cardiac fibroblasts under identical treatment (Fig.1N and Fig.S1H).

To investigate the expression and distribution of ZDHHC14 in the human heart, we analyzed data from the Human Protein Atlas (https://www.proteinatlas.org/). Collectively, these findings demonstrate that ZDHHC14 is present in multiple resident cardiac cell populations, with notably higher abundance observed in cardiomyocytes (Figures S1I-J). Enhanced ZDHHC14 immunostaining was evident in TAC-operated myocardial tissues (Fig.1O, Fig.S1K). Taken together, the data indicate that ZDHHC14 is upregulated in hypertrophic cardiac tissue from both murine and human sources.

### Cardiac-specific knockdown of ZDHHC14 suppress myocardial hypertrophy in vitro and in vivo

To manipulate ZDHHC14 expression in NRCMs, we employed adenoviral and siRNA transfection, with successful overexpression and knockdown confirmed via immunoblotting (Fig.S2A-B). In AngII stimulated NRCMs, α-actin immunofluorescence revealed that ZDHHC14 knockdown markedly reduced cell surface area relative to control (Fig.2A). Quantitative analysis further demonstrated that ZDHHC14 silencing blunted the AngII-evoked rise in atrial natriuretic peptide (ANP) and myosin heavy chain 7 (MYH7) mRNA levels (Fig.2B), whereas ZDHHC14 gain- of-function reversed this suppressive effect (Fig. S2C-D). For in vivo manipulation, we constructed an AAV9 vector carrying a cTNT promoter-driven shRNA against ZDHHC14 (AAV9-shZDHHC14) to achieve cardiomyocyte-restricted gene silencing. Following a two-week pre-treatment period, animals were subjected to TAC surgery to establish pressure overload-induced hypertrophy (experimental timeline depicted in Fig.2C). Immunoblot analysis verified efficient ZDHHC14 knockdown in mice cardiac tissue (Fig.S2E). ZDHHC14 deficiency significantly ameliorated TAC-evoked deterioration of cardiac function (Fig.2D-F). Moreover, histological assessments showed that ZDHHC14 knockdown mitigated myocyte enlargement, evidenced by hematoxylin and eosin (H&E) staining (Fig.2G), decreased heart weight-to-body weight ratios (HW/BW, Fig. 2H) and heart weight-to-tibial length ratios (HW/TL, Fig. 2I), as well as wheat germ agglutinin (WGA) staining results (Fig. 2J-K). Masson’s trichrome staining indicated attenuated cardiac fibrosis following ZDHHC14 knockdown (Fig. 2L-M), and RT-qPCR confirmed reduced mRNA levels of ANP and MYH7 in ZDHHC14 knockdown hearts (Fig. 2N). In aggregate, these findings establish that ZDHHC14 knockdown confers protection against pressure overload-induced cardiac dysfunction and hypertrophic remodeling.

**Fig. 2.**
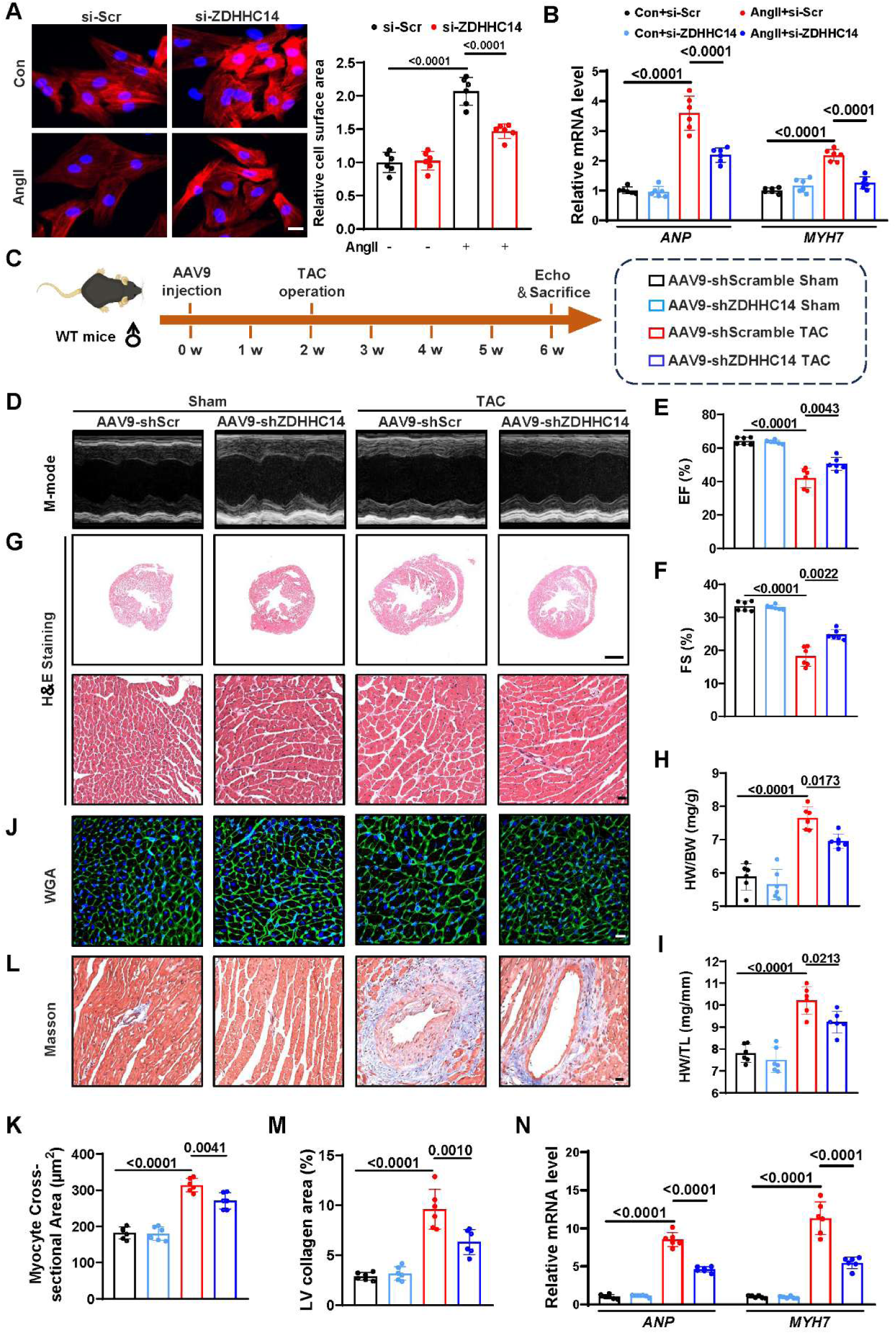
ZDHHC14 cardiomyocyte-specific knockdown inhibits TAC induced cardiac hypertrophy. A. NRCMs were transfected with scramble control siRNA (si- Scr) or ZDHHC14 siRNA (si-ZDHHC14) and then cultured in the presence or absence of Ang II (1 μM) for 24 h. To verify the presence of cardiomyocytes, we performed α-Actinin staining. Representative images (left panel) and statistical results (right panel) of NRCM cell size (n = 6 biological replicates per group). Scale bar: 20 µm. B. NRCMs were treated as shown in (A), and then we performed qRT-PCR (B) to determine the ANP and MYH7 mRNA levels (n = 6 biological replicates per group). C. Experimental timeline: mice were injected with the AAV9-cTNT-ZDHHC14-shRNA (AAV9-sh ZDHHC14) or the scramble control (AAV9-shScr; 2×10^11^ particles per mouse) via the tail vein for 2 weeks; subsequently, these mice underwent TAC surgery and were examined 4 weeks later. D-F. Representative M-mode echocardiography of left ventricular chamber and measurement of EF%, FS% (n = 6 biological replicates per group). G. Representative HE staining images of cardiac sections (n = 6 biological replicates per group). Scale bar, 1 mm and 25 μm. H. The ratio of HW to BW (n = 6 biological replicates per group). I. The ratio of HW to TL (n = 6 biological replicates per group). J. Representative WGA staining images (n = 6 biological replicates per group). Scale bar, 20 μm. K. Quantitative analysis of WGA staining images for heart sections (n = 6 biological replicates per group). L-M. Representative masson stained images in heart sections (L) and quantitative analysis (M) (n = 6 biological replicates per group). Scale bar, 20 μm. N. Relative mRNA level of ANP and MYH7 in the heart tissues (n = 6 biological replicates per group). Abbreviations: si: silence; Scr: scramble; Echo: Echocardiography; EF: Ejection Fraction; FS: Fractional Shortening; LV: left vetricular. For statistical analysis, two-way ANOVA followed by Tukey post hoc tests was used for (A, B, E, F, H, I, K, M, and N). Data are presented as mean ± SEM. P < 0.05 indicates a statistically significant difference.

### Cardiac-specific overexpression of ZDHHC14 aggravates myocardial hypertrophy

To probe whether ZDHHC14 exerts a pathogenic role in cardiac hypertrophy, we used an AAV9-cTNT vector to elevate ZDHHC14 levels specifically in mice cardiomyocytes (Fig.3A), with overexpression confirmed by western blot analysis (Fig.S3A). Viral injection was followed two weeks later by TAC surgery. ZDHHC14-overexpressing hearts displayed pronounced exacerbation of both systolic dysfunction and hypertrophic manifestations at 4 weeks after TAC (Fig.3B-D). Histological measurements revealed that ZDHHC14 overexpression promotes myocyte enlargement, evidenced by H&E staining (Fig.3E), increased HW/BW ratio (Fig.3F) and HW/TL ratio (Fig.3G). Microscopic assessments further demonstrated that ZDHHC14 overexpression expanded myocyte cross-sectional area (Fig.3H-I), intensified cardiac fibrosis (Fig.3J-K), and drove upregulation of the hypertrophy-associated genes ANP and MYH7 (Fig.3L). These observations collectively support the conclusion that cardiomyocyte ZDHHC14 overexpression is sufficient to worsen pressure overload-induced cardiac pathological progression.

**Fig. 3.**
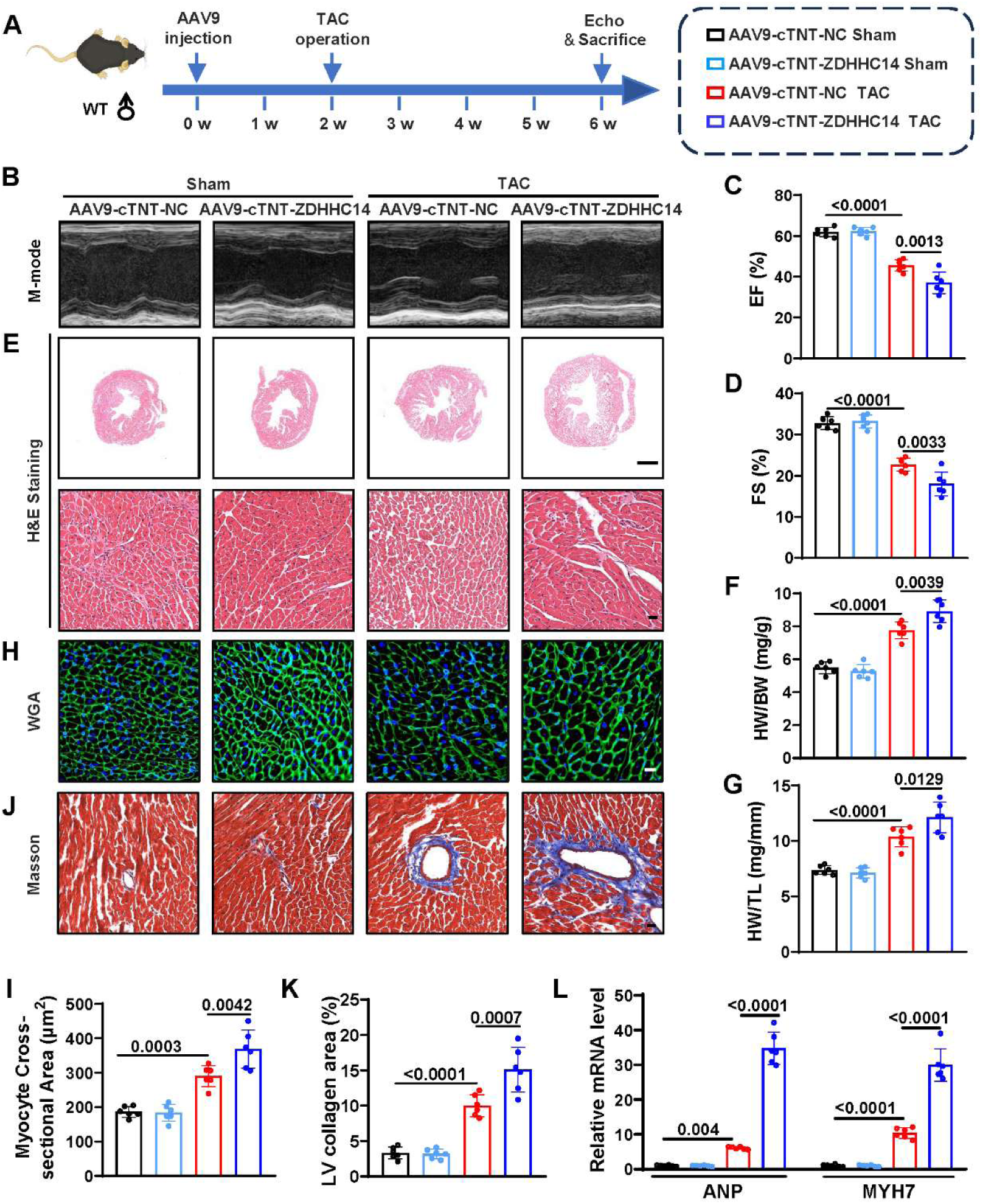
Cardiomyocyte specific overexpression of ZDHHC14 aggravates TAC induced cardiac dysfunction and hypertrophy. A. Experimental timeline: mice were injected with the AAV9-cTNT-ZDHHC14 or the negative control (AAV9-cTNT-NC; 2 ×10^11^ particles per mouse) via the tail vein for 2 weeks; subsequently, these mice underwent TAC surgery and were examined 4 weeks later. B-D. Representative M-mode echocardiography of left ventricular chamber and measurement of EF%, and FS% (n = 6 biological replicates per group). E. Representative HE staining images of cardiac sections (n = 6 biological replicates). Scale bar, 1 mm and 25 μm. F. The ratio of HW to BW (n = 6 biological replicates per group). G. The ratio of HW to TL (n = 6 biological replicates per group). H. Representative WGA staining images (n = 6 biological replicates per group). Scale bar, 20 μm. I. Quantitative analysis of WGA staining images for heart sections (n = 6 biological replicates per group). J-K. Representative masson stained images in heart sections (J) and quantitative analysis (K) (n = 6 biological replicates per group). Scale bar, 20 μm. L. Relative mRNA level of ANP and MYH7 in the heart tissues (n = 6 biological replicates per group). For statistical analysis, two-way ANOVA followed by Tukey post hoc tests was used for (C, D, F, G, I, K and L). Data are presented as mean ± SEM. P < 0.05 indicates a statistically significant difference.

### ZDHHC14 promotes cardiac hypertrophy through activation of IFN-I signaling

To elucidate the molecular underpinnings of ZDHHC14 in hypertrophic cardiac remodeling, we conducted RNA-sequencing on Ang II-stimulated NRCMs following transfection with either control siRNA or ZDHHC14-targeting siRNA. Differential expression analysis, visualized as a volcano plot, identified 357 transcripts that were significantly upregulated and 792 that were downregulated (Figure 4A). Gene Ontology (GO) enrichment analysis highlighted DEGs were enriched in inflammatory response regulation and IFN-I signaling (Figure 4B). Furthermore, gene set enrichment analysis (GSEA) corroborated these findings, demonstrating substantial enrichment of the IFN-I signaling pathway among the differentially expressed gene sets (Figure 4C). Likewise, GSEA showed significant enrichment of the DEGs in the cytosolic DNA-sensing pathway (Fig.S4A), a pathway intimately linked to IFN-I signaling. Given that previous investigations have established a critical role for IFN-I signaling in pathological cardiac hypertrophy^9^, we next examined whether ZDHHC14 modulates this signaling. We began by examining whether ZDHHC14 modulates IFN-I signaling in vivo. At 4 weeks following TAC, hearts with ZDHHC14 knockdown exhibited marked suppression of the IFN-I signaling, as evidenced by reduced phosphorylation levels of STING, TBK1, and IRF3, alongside diminished mRNA levels of representative type I interferon genes (Ifna and Ifnb1) (Figure 4D-H). To complement these observations in vitro, we conducted both loss- and gain-of-function experiments in NRCMs. RT-qPCR and immunoblot analyses revealed that Ang II treatment enhanced the activation of IFN-I signaling, which was effectively blunted by ZDHHC14 silencing (Fig.4I-M). Moreover, ZDHHC14 overexpression further potentiated IFN-I signaling (Figure 4N-P and Fig.S4B-C). Collectively, these data demonstrated that ZDHHC14 acts as a positive regulator of IFN-I signaling activation in the setting of cardiac hypertrophy. GSK8612 is known to inhibit IFN-I signaling by targeting TBK1 phosphorylation^31^. When we applied this compound to ZDHHC14-overexpressing NRCMs, it significantly reversed the enlargement of cell size and the upregulation of hypertrophy-related genes triggered by ZDHHC14 overexpression (Fig.4Q-S). Collectively, these data indicate that ZDHHC14 promotes pathological cardiac hypertrophy via IFN-I signaling activation.

**Fig. 4.**
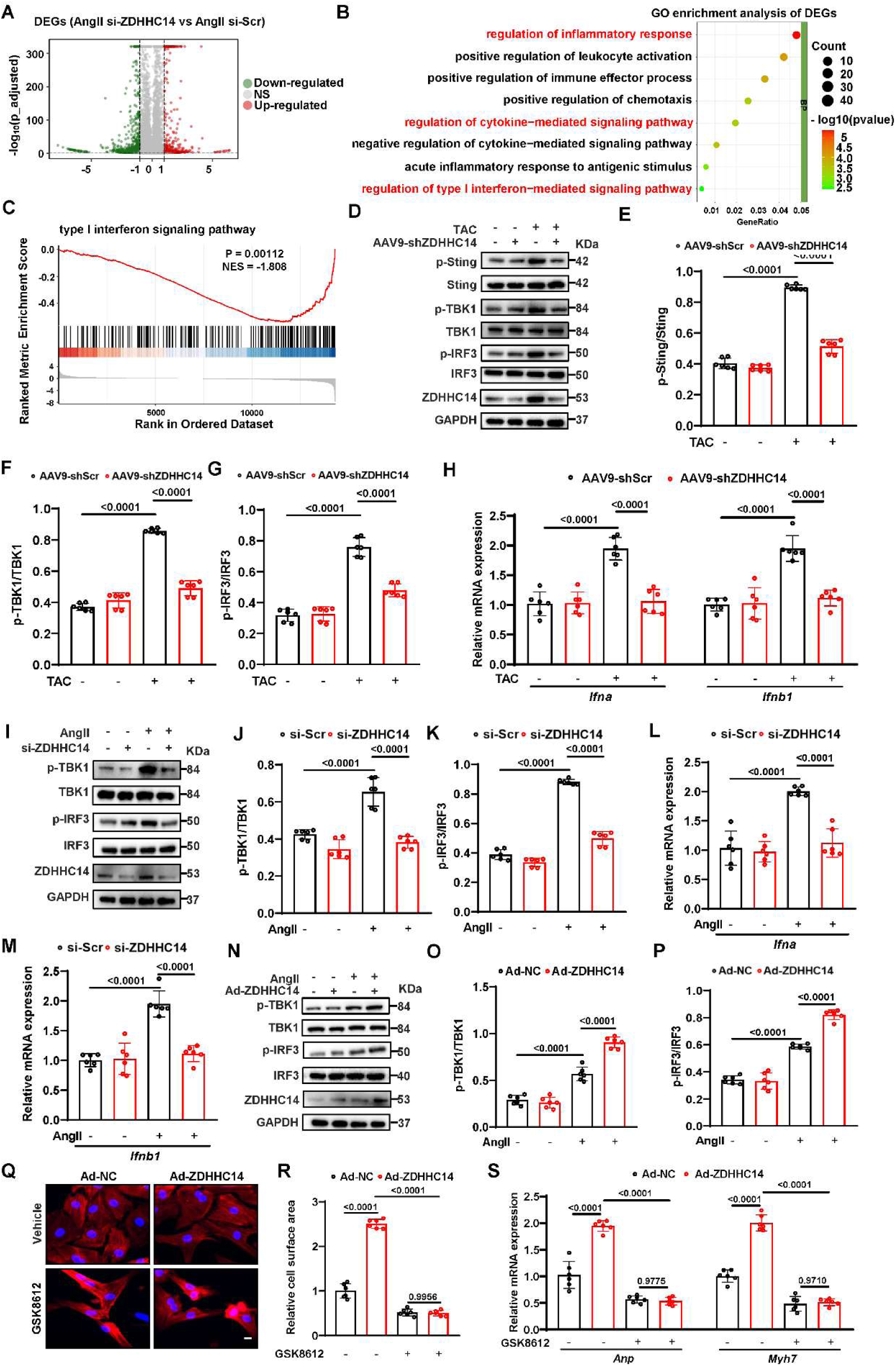
ZDHHC14 promotes cardiac hypertrophy through activation of type I interferon (IFN-I) signaling. A. Volcano plot of DEGs between si-ZDHHC14 and si- Scr group in NRCMs (n = 3 biological replicates per group). B. GO pathway enrichment analysis of the identified DEGs based on RNA-seq data set from Ang II-treated NRCMs infected by control or ZDHHC14 siRNA (n = 3 biological replicates per group). C. GSEA of DEGs for type I interferon (IFN-I) signaling. D-G. Western blots of total and phosphorylated levels of IFN-I signaling-related proteins from heart tissues of mice injected with the AAV9-shZDHHC14 or AAV9-shScr after the sham or TAC surgery (n = 6 biological replicates per group). H. The mRNA levels of representative type I interferon genes (Ifna and Ifnb1) from heart tissues of mice injected with the AAV9-shZDHHC14 or AAV9-shScr after the sham or TAC surgery (n = 6 biological replicates per group). I-K. Western blots of total and phosphorylated levels of IFN-I signaling-related proteins at 24 h after PBS or Ang II (1 μM) treatment in NRCMs infected with si-ZDHHC14. si-Scr used as a control (n = 6 biological replicates per group). L-M. The mRNA levels of representative type I interferon genes at 24 h after PBS or Ang II (1 μM) treatment in NRCMs infected with si-ZDHHC14. si-Scr used as a control (n = 6 biological replicates per group). N-P. Western blots of total and phosphorylated levels of IFN-I signaling-related proteins at 24 h after PBS or Ang II (1 μM) treatment in NRCMs infected with Ad-ZDHHC14. Ad-NC used as a control (n = 6 biological replicates per group). Q-R. Representative images (left) and quantitative results of the cell surface area (right) in NRCMs infected with the indicated adenovirus and treated with or without GSK8612 (80 μM) for 24 h (n = 6 biological replicates). Scale bar: 20 µm. S. The ANP and MYH7 mRNA expression levels in NRCMs infected with the indicated adenovirus and treated with or without GSK8612 (80 μM) for 24 h (n = 6 biological replicates). For statistical analysis, two-way ANOVA followed by Tukey post hoc tests was used for (E-H, J-M, O-P and R-S). Data are presented as mean ± SEM. P < 0.05 indicates a statistically significant difference.

### Identification of TBK1 as a potential substrate of ZDHHC14

We next sought to elucidate how ZDHHC14 potentiates IFN-I signaling. Since PATs typically act through substrate-specific interactions^32^, we mapped the ZDHHC14 interactome by performing immunoprecipitation (IP) coupled with liquid chromatography–tandem mass spectrometry (LC–MS/MS) in HEK 293T cells overexpressing ZDHHC14. In total, 1,138 proteins were identified as potential substrates of ZDHHC14 by this screening (Fig.5A-B). Ranking these candidates by unique peptide counts revealed that TBK1 exhibited the highest score (Fig.5C and Fig.S5A). TBK1 is a critical node in the IFN-I pathway, governing IRF3/interferon regulatory factor 7 (IRF7) phosphorylation and nuclear translocation to initiate downstream interferon gene transcription^33^. Molecular docking further supported a robust interaction between ZDHHC14 and TBK1 (Fig.5D). Co-IP assays in NRCMs confirmed the co-precipitation of ZDHHC14 with TBK1 (Fig.5E-F), and immunofluorescence staining demonstrated their overlapping subcellular distribution (Fig.5G). Independent cotransfection of Flag-ZDHHC14 and HA-TBK1 into HEK 293T cells corroborated this binding (Fig.5H-I). Notably, AngII treatment significantly enhanced the association between ZDHHC14 and TBK1 in NRCMs (Fig.S5B). TBK1 comprises three conserved regions: a kinase domain (KD), a ubiquitin-like domain (ULD), and a coiled-coil (CC) domain^34^. To map the palmitoylation site on TBK1, we transfected HEK 293T cells with constructs encoding full-length (FL) TBK1, the KD alone, or the ULD-CC tandem. The results showed that ZDHHC14 specifically binds with the TBK1 KD region (Fig. 5J). These results confirm the existence of a direct interaction between ZDHHC14 and TBK1.

**Fig. 5.**
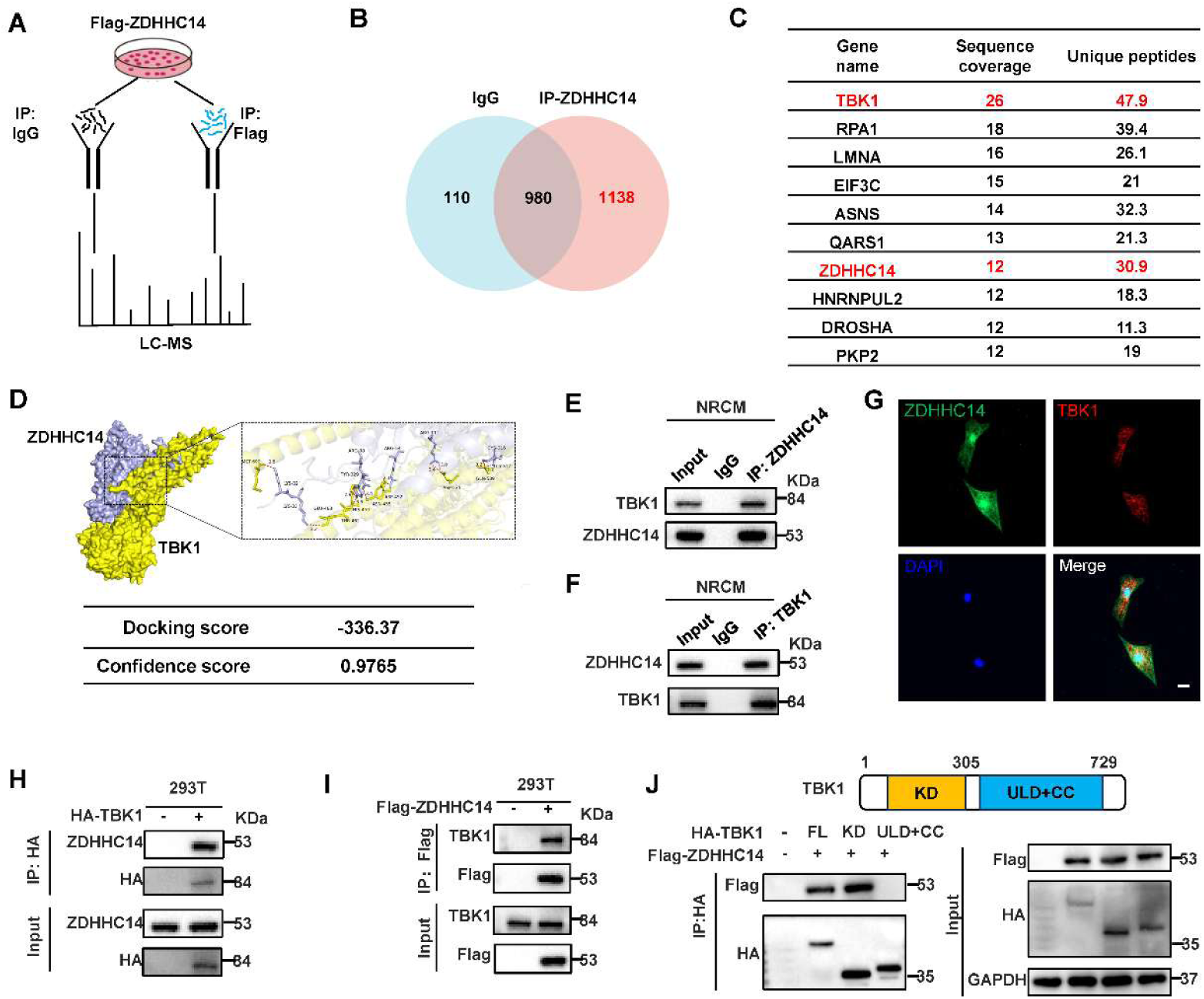
Identification of TBK1 as a potential substrate of ZDHHC14. A. Schematic diagram showing the MS (mass spectrometry) analysis to identify the specific target of ZDHHC14. B. The venn diagram shows potential ZDHHC14 substrates identified from the interactomes. C. The list is the top 10 potential ZDHHC14 substrates sorted by unique peptide. D. The molecular docking between ZDHHC14 and TBK1 constructed by HDOCK software. E-F. Immunoprecipitation (IP) was performed in cardiomyocyte lysate from neonatal rat cardiomyocytes (NRCMs) with rabbit IgG, anti-ZDHHC14, or anti-TBK1 antibodies, and ZDHHC14 and TBK1 were detected by Western blot analysis (n = 6 biological replicates). G. Immunofluorescence assays verified the binding of ZDHHC14 with TBK1 in NRCMs (n = 6 biological replicates). Scale bar, 50 μm. H-I. Representative western blots of Co-IP assays in HEK 293T cells transfected with Flag-tagged ZDHHC14 and HA-tagged TBK1. Flag and HA antibodies were used as western blot probes (n = 6 biological replicates). J. Co-IP showed that the KD region of TBK1 is essential for the interaction with ZDHHC14 and TBK1(n = 3 biological replicates).

### ZDHHC14 potentiates IFN-I signaling by promoting TBK1 palmitoylation and **subsequent phosphorylation activation**

To determine whether TBK1 undergoes palmitoyl modification, we employed an acyl-biotin exchange (ABE) assay. Retention of the chemiluminescent signal following hydroxylamine (HAM) treatment confirmed the presence of thioester-linked palmitate groups on the protein^35^. Our data showed that TBK1 is indeed palmitoylated (Fig. 6A). We next observed that ZDHHC14 knockdown substantially reduced TBK1 palmitoylation (Fig.6B), whereas ZDHHC14 overexpression promoted TBK1 palmitoylation (Fig. 6C). Immunoblot analysis further revealed that both the full-length TBK1 (FL) and the isolated KD region were palmitoylated, whereas the ULD-CC fragment was not (Fig.6D). Using the GPS-Palm tool^36^, we identified five top candidate palmitoylation residues on TBK1(C89, C91, C267, C292, and C728) (Fig. 6E), all of which are highly conserved across vertebrate species (Fig. 6F). Among these, C89, C91, C267, and C292 reside within the KD region. Site-directed mutagenesis demonstrated that only the C267S mutant drastically diminished TBK1 palmitoylation (Fig.6G), pinpointing C267 as the principal palmitoylation site. Furthermore, TBK1 C267S mutation abrogated ZDHHC14-driven phosphorylation of both TBK1 and IRF3, and concurrently blunted the upregulation of type I interferon genes (Fig.6H-L). Likewise, disrupting the catalytic cysteine (C195S) of ZDHHC14 abolished its ability to potentiate IFN-I signaling (Fig.6M-O and Fig.S6A). Collectively, these findings establish that ZDHHC14 mediated palmitoylation of TBK1 at C267 site is critical for TBK1 phosphorylation and the activation of the type I interferon signaling.

**Fig. 6.**
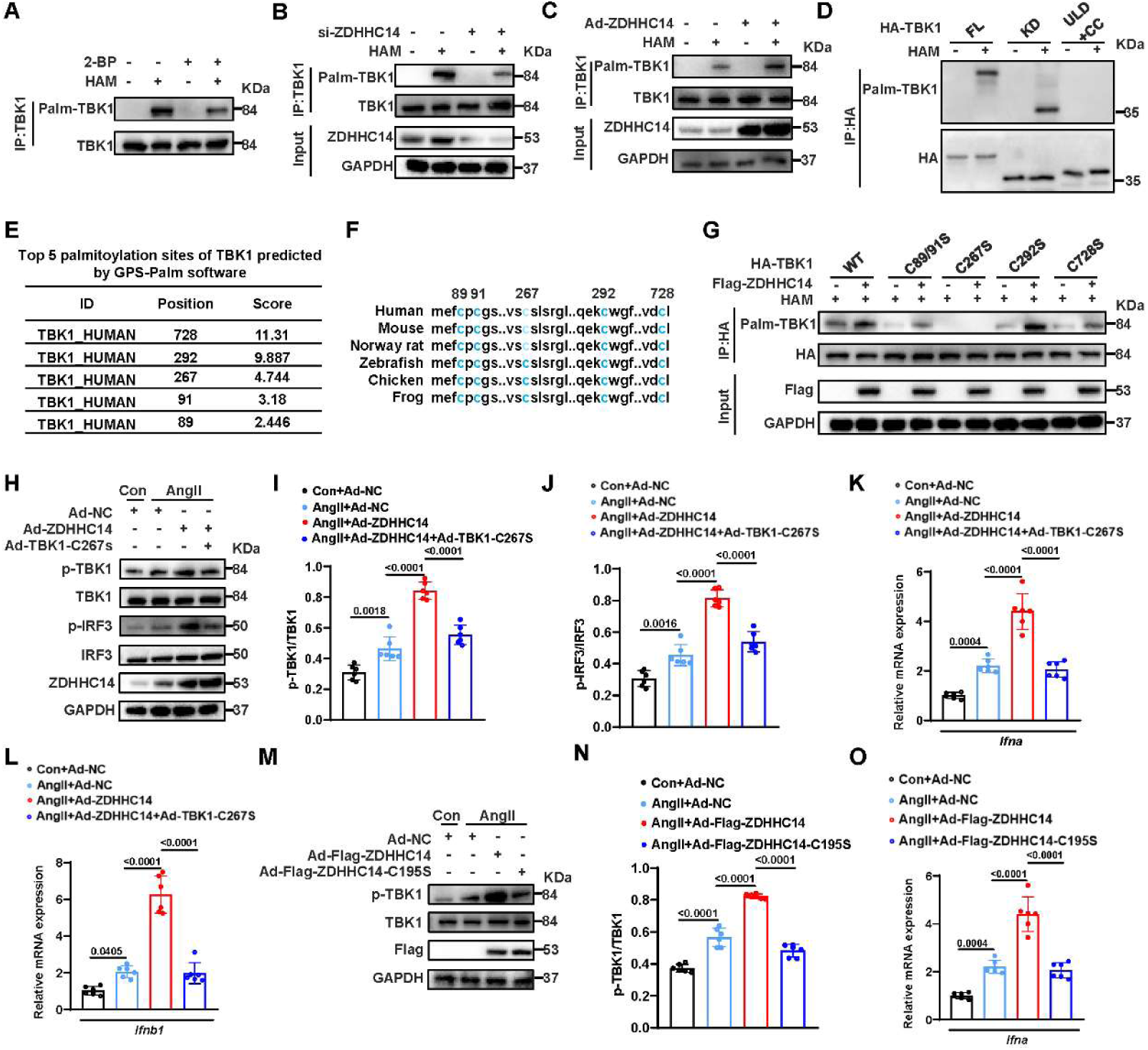
ZDHHC14 potentiates IFN-I signaling by promoting TBK1 palmitoylation and subsequent phosphorylation activation. A. TBK1 palmitoylation levels were measured using ABE assay and immunoblotting (n = 6 biological replicates). B. NRCMs were transfected with scramble control siRNA (si-Scr) or ZDHHC14 siRNA (si-ZDHHC14) for 24 h. And then ABE assay was used to measure TBK1 palmitoylation level (n = 6 biological replicates). C. NRCMs were transfected with negative control (Ad-NC) or ZDHHC14 overexpression adenoviruses (Ad- ZDHHC14) for 24 h. And then ABE assay was used to measure TBK1 palmitoylation level (n = 6 biological replicates). D. HEK293T cells were transfected with wild-type (WT) or deletion mutants of HA-TBK1 for 24 h, and TBK1 palmitoylation levels were measured using ABE assay and immunoblotting (n = 6 biological replicates). E. Top 5 palmitoylation sites of human TBK1 predicted by GPS-Palm software. F. Comparative analysis of potential palmitoylation sites within TBK1 sequences across different species. G. HEK 293T cells were transfected with the vector plasmid or Flag-ZDHHC14, and WT HA-TBK1 or TBK1 palmitoylation-deficient mutants for 24 h. TBK1 palmitoylation levels were detected by ABE assay and immunoblot analysis (n = 6 biological replicates). H-J. Western blot analyses of p-TBK1 and p-IRF3 levels with overexpression of the indicated groups in NRCMs under Ang II (1 μM) stimulation for 24 hours (n = 6 biological replicates). K-L. The mRNA levels of IFN-I genes in indicated group (n = 6 biological replicates). M-N. Western blot analyses of p-TBK1 and TBK1 in the indicated groups (n = 6 biological replicates). O. The mRNA levels of Ifna in indicated group (n = 6 biological replicates). For statistical analysis, one-way ANOVA/Tukey test was used for (I-L, N and O). Data are presented as mean ± SEM. P < 0.05 indicates a statistically significant difference.

### Palmitoylation of TBK1 at C267 site Mediates ZDHHC14-induced Cardiac Dysfunction and Hypertrophy

To further explore the functional significance of TBK1 C267 site palmitoylation in ZDHHC14-induced cardiac hypertrophy, we examined the effect of TBK1 C267 mutation in NRCMs. As shown in Fig.7A-B, TBK1 C267S mutant significantly reversed the pro-hypertrophic effects induced by ZDHHC14 overexpression in vitro. To further confirm whether TBK1 C267 palmitoylation in cardiomyocytes contributes to ZDHHC14-induced cardiac hypertrophy in vivo, we delivered cardiomyocyte-specific AAV9 encoding either TBK1-WT or TBK1-C267S mutation via tail vein injection into mice with cardiac-specific ZDHHC14 overexpression (experimental timeline shown in Fig.7C). Remarkably, mice injected with TBK1 C267 mutation adenovirus substantially reversed ZDHHC14-driven cardiac dysfunction (Fig.7D-F). Furthermore, mice injected with TBK1 C267 mutation adenovirus prevented the pathological increases in heart dimensions (Fig.7G), HW/BW ratio (Fig.7H), HW/TL ratio (Fig.7I), and cardiomyocyte area (Fig. 6J-K) in ZDHHC14 overexpression mice. mice injected with TBK1 C267 mutation adenovirus also ameliorated myocardial fibrosis (Fig.7L-M) and normalized the elevated expression of hypertrophy markers (Anp and Myh7) caused by ZDHHC14 overexpression (Fig.7N). Furthermore, mice injected with TBK1 C267 mutation adenovirus abrogated ZDHHC14-driven phosphorylation of STING, TBK1, and IRF3(Fig.S7A). Collectively, these results establish that palmitoylation of TBK1 at C267 serves as a critical downstream effector through which ZDHHC14 modulates hypertrophic remodeling in response to pressure overload.

**Fig. 7.**
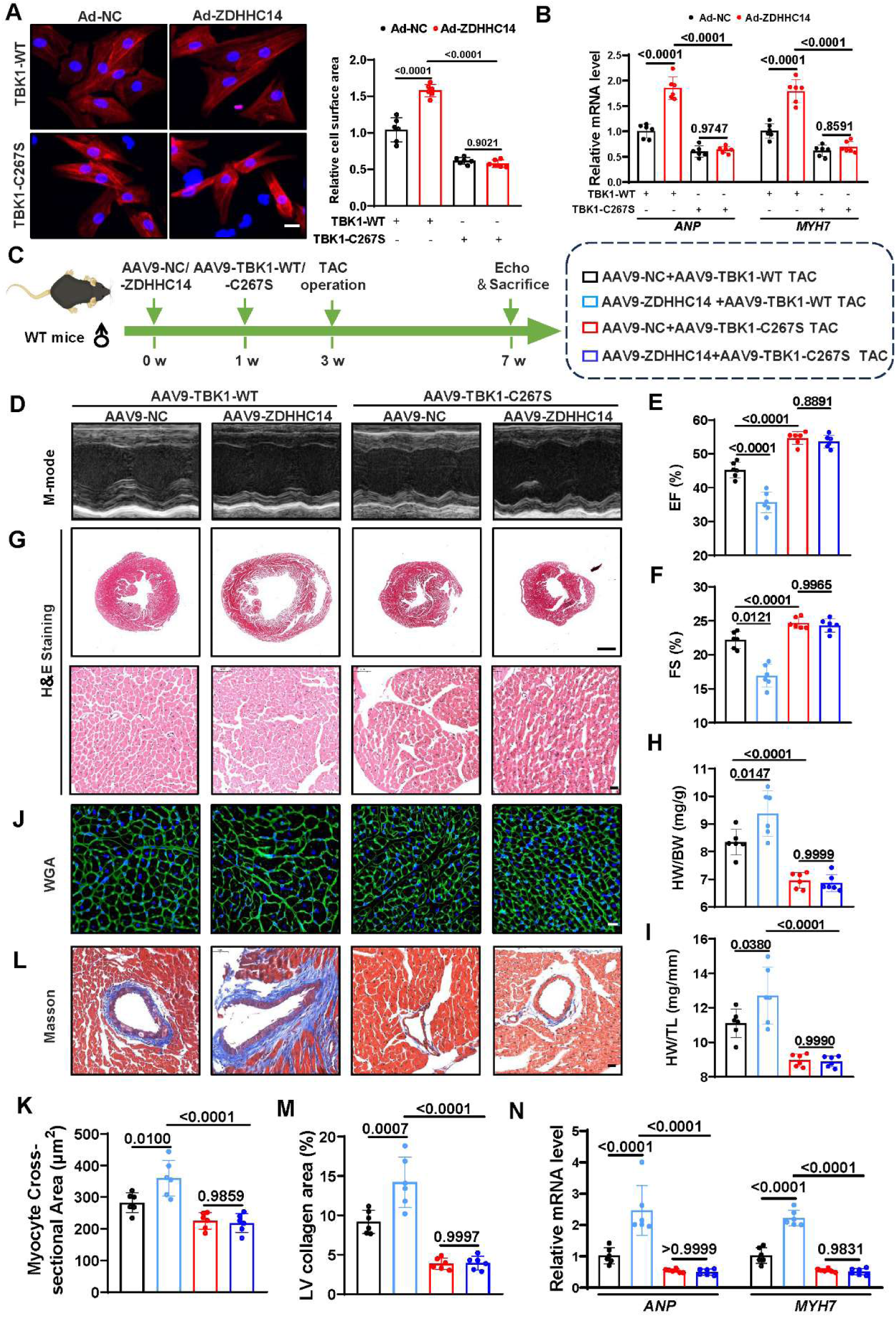
Palmitoylation of TBK1 at C267 site Mediates ZDHHC14-induced Cardiac Dysfunction and Hypertrophy. A. Representative immunofluorescence images (left) and quantitative results of the cell surface area (right) of NRCMs infected with the indicated adenovirus and treated with Ang II (1 μM) for 24 h (n = 6 biological replicates). Scale bar, 20 μm. B. The ANP and MYH7 mRNA expression levels of NRCMs infected with the indicated adenovirus and treated with Ang II (1 μM) for 24 h (n = 6 biological replicates). C. Experimental timeline: male wildtype mice were injected with the AAV9-cTNT-OTUD5 or negative control (AAV9-cTNT-NC; 1×1011 particles per mouse) via the tail vein for 1 week; subsequently, these mice were injected with the AAV9-cTNT-TBK1-C267S or the wildtype TBK1 (AAV9-cTNT-TBK1-WT; 1×10^11^ particles per mouse) via the tail vein for 2 weeks. Then these mice underwent TAC surgery and were examined 4 weeks later. D-F. Representative M-mode echocardiography of left ventricular chamber and measurement of EF%, and FS% in the indicated groups (n = 6 biological replicates per group). G. Representative HE staining images of cardiac sections in the indicated groups (n = 6 biological replicates per group). Scale bar, 0.5 mm and 20 μm. H. The ratio of HW to BW in the indicated groups (n = 6 biological replicates per group). I. The ratio of HW to TL in the indicated groups (n = 6 biological replicates per group). J. Representative WGA staining images in the indicated groups (n = 6 biological replicates per group). Scale bar, 20 μm. K. Quantitative analysis of WGA staining images for heart sections in the indicated groups (n = 6 biological replicates per group). L-M. Representative masson stained images of myocardial interstitium in heart sections in the indicated groups (n = 6 biological replicates per group). Scale bar, 50 μm (L) and quantitative analysis (M). N. The mRNA level of ANP and MYH7 in the heart tissues in the indicated groups (n = 6 biological replicates per group). For statistical analysis, two-way ANOVA followed by Tukey post hoc tests was used for (A, B, E, F, H, I, K, M, and N). Data are presented as mean ± SEM. P < 0.05 indicates a statistically significant difference.

## Discussion

In this study, elevated ZDHHC14 levels were consistently detected in hypertrophic cardiac tissue from both clinical samples and experimental mice. Cardiomyocyte-restricted knockdown of ZDHHC14 protected against TAC-induced heart failure and hypertrophy, whereas its overexpression aggravated these pathological features. At the mechanistic level, ZDHHC14 engages TBK1 in a physical complex and drives its palmitoylation at C267 site, leading to increased TBK1 phospho-activation. This ZDHHC14-TBK1 axis subsequently instigates IFN-I signaling, which promotes the progression of cardiac hypertrophy.

S-palmitoylation, defined as the reversible cysteine-linked conjugation of palmitate (C16:0), modulates protein amphiphilicity and mediates recruitment to lipid-raft platforms, where it exerts control over receptor recycling, signaling-fidelity, and scaffold-protein assembly^30^. This regulatory lipid switch has profound implications for cardiovascular homeostasis: in endothelial cells, its misregulation attenuates NO-mediated vasodilatory capacity and instigates a pro-adhesive, pro-oxidative milieu that seeds atherogenic lesions^37, 38^; in cardiomyocytes, it calibrates mechanical performance, ionic conductance, and ER-stress tolerance, such that derangements in palmitoylation favor apoptotic attrition and interstitial fibrosis, thereby perpetuating maladaptive remodeling and worsening hemodynamic failure^39, 40^. Inflammation plays a pivotal role in pathological cardiac hypertrophy and heart failure^41^. However, the involvement of S-palmitoylation in this pathological process has not been previously reported. Our study reveals a critical mechanism by which ZDHHC14-mediated S-palmitoylation of TBK1 drives IFN-I signaling and inflammatory activation, thereby filling a gap in our understanding of pathological cardiac hypertrophy.

As a critical member of the palmitoyl acyltransferase family, ZDHHC14 regulates the palmitoylation of diverse substrate proteins, thereby influencing a broad spectrum of biological processes—including ferroptosis^26^, cell cycle progression^27^, metabolism^28^, and ion channel function^29^—and consequently promoting the progression of both tumors and neurological disorders. In recent years, emerging evidence has implicated ZDHHC14 in inflammation-related pathologies^42, 43^. For instance, ZDHHC14-mediated TEA Domain Transcription Factor 4 (TEAD4) palmitoylation drives Th17 cell recruitment in renal immunopathology, exacerbating inflammatory injury^42^. Likewise, polystyrene nanoplastics (PS-NPs) have been shown to upregulate ZDHHC14, which in turn promotes Solute Carrier Family 31 Member 1 (SLC31A1) palmitoylation in macrophages, leading to NLR Family Pyrin Domain Containing 3 (NLRP3) pathway activation in alveolar epithelial cells^43^. Building upon this context, our study is the first to demonstrate a positive regulatory role of ZDHHC14 in inflammatory activation and IFN-I signaling within the setting of pathological cardiac hypertrophy.

As a critical member of the IFN-I family, the phosphorylation and activation of TBK1 play a pivotal role in pathological cardiac hypertrophy^12^. Accumulating evidence has revealed that TBK1 kinase activity is finely tuned by various other PTMs, including ubiquitination^12^, acetylation^44^, and methylation^45^, which can directly or indirectly modulate its phosphorylation-dependent activation. In this study, we identify for the first time that ZDHHC14-mediated S-palmitoylation of TBK1 promotes its phosphorylation and activation; however, the precise molecular mechanisms underlying this regulatory axis warrant further investigation.

Our study identifies a novel role for cardiomyocyte-specific ZDHHC14 overexpression in exacerbating TAC-induced cardiac fibrosis, a finding that has not been previously reported. Although the direct link between ZDHHC14 and cardiac fibrosis remains unexplored, our mechanistic investigations, together with existing evidence, provide a plausible framework to elucidate this regulatory process. We demonstrate that ZDHHC14 promotes cardiac hypertrophy by enhancing TBK1 palmitoylation and activating IFN-I signaling. Notably, TBK1 has been well established as a key mediator of both cardiac hypertrophy and fibrosis^46^, largely through its ability to activate the NF-κB signaling pathway. Collectively, our study uncovers a novel function of ZDHHC14 in promoting cardiac fibrosis via TBK1-dependent activation. These findings not only identify ZDHHC14 as a potential therapeutic target for fibrotic heart diseases but also offer new insights into the intricate interplay between palmitoylation, inflammatory signaling, and cardiac remodeling.

Admittedly, there are certain limitations in this study that warrant further refinement. Aside from the ZDHHC14-TBK1 axis, ZDHHC14 has been reported to deubiquitinate other substrates. we cannot entirely rule out the possibility that ZDHHC14 regulates the hypertrophic response of cardiomyocytes through other substrates. Despite the intriguing findings of this study, we acknowledge a notable limitation regarding the clinical sample size. The current analysis was based on only 8 patients with cardiac hypertrophy and 4 healthy controls. This relatively small sample size may restrict the statistical power of our results. In addition, we identify that ZDHHC14-mediated S-palmitoylation of TBK1 promotes its phosphorylation and activation; however, the precise molecular mechanisms underlying this regulatory axis warrant further investigation.

Collectively, our results demonstrate that ZDHHC14 associates with TBK1 to form a physical complex and drives its palmitoylation at residue C267, leading to enhanced TBK1 phosphorylation and subsequent activation of IFN-I signaling, thereby promoting the progression of pathological cardiac hypertrophy. This discovery expands the current knowledge regarding the functional contributions of palmitoyl acyltransferases in cardiac hypertrophy. Additionally, the potential of ZDHHC14 inhibition as a therapeutic approach for cardiac hypertrophy warrants further exploration.

## Methods

### Human heart sample

The human cardiac tissue samples employed in this study were obtained from eight patients diagnosed with cardiac hypertrophy (CH) who underwent artificial heart implantation, as well as four non-cardiac hypertrophy (NCH) subjects who experienced accidental fatalities and were admitted to the First Affiliated Hospital of Nanjing Medical University. Clinical details of the participants are provided in Table S1. Written informed consent was secured from all patients prior to sample collection. Ethical approval for this research was issued by the Ethics Committee of the First Affiliated Hospital of Nanjing Medical University (approval number: 2025-SR-718).

### Animals and treatments

All animal procedures were conducted in strict compliance with the National Institutes of Health (NIH) guidelines (Publication No. 85-23, revised 2011) and were approved by the Institutional Animal Care and Use Committee of Yangzhou University (Approval No. 202607027). Male wild-type (WT) C57BL/6 mice, acquired from GemPharmatech (Nanjing, China), were acclimatized to the laboratory environment for one week prior to experimentation. Animals were housed under specific pathogen-free conditions at 20–25°C with 50 ± 5% humidity and had unrestricted access to standard chow and autoclaved water. To establish pressure overload-induced cardiac remodeling, animals underwent transverse aortic constriction (TAC) for a duration of four weeks, as described previously^47^. For angiotensin II (Ang II)-induced hypertrophy, twelve healthy male mice (6-8 weeks old) received either Ang II (1 μg/kg/min) or an equivalent volume of normal saline via subcutaneously implanted osmotic minipumps (Alzet MODEL 1004, USA) over four weeks (n = 6 per group). For in vivo functional assessment of ZDHHC14, mice were pretreated with tail-vein injections of adeno-associated virus 9 (AAV9)-cTNT-ZDHHC14 or AAV9-cTNT-NC (negative control) two weeks prior to TAC or sham surgery to achieve cardiac-specific overexpression; conversely, AAV9-shZDHHC14 or AAV9-shScramble (shScr) were administered to suppress ZDHHC14 expression. Each animal received 1×10^11^ viral genome copies in 100 µL phosphate-buffered saline (PBS) at six weeks of age. To evaluate the contribution of TBK1 C267 palmitoylation, additional cohorts of WT male mice received AAV9-TBK1-C267S or AAV9-TBK1-WT virus two weeks before TAC or sham operations. All viral constructs were sourced from Genepharma (Shanghai, China). Echocardiographic assessment of cardiac performance was performed using a Vevo 3100 high-resolution imaging system. At the experimental endpoint, mice were euthanized humanely under pentobarbital sodium anesthesia, and myocardial specimens were harvested for further analyses.

### Statistical analysis

All quantitative results are presented as mean values ± standard error of the mean (SEM). Intergroup disparities between two experimental cohorts were evaluated using Student’s unpaired t-test. For single-factor comparisons across three or more groups, we applied one-way analysis of variance (ANOVA); for multifactorial designs involving two independent variables, two-way ANOVA was performed. All statistical computations were executed with GraphPad Prism 8.0 (GraphPad Software, San Diego, CA, USA). A significance threshold of P < 0.05 was adopted for all hypothesis tests. Comprehensive procedural descriptions are accessible in the Supplemental Materials.

## Acknowledgments

We wish to extend our heartfelt thanks to everyone who contributed their support throughout the creation of this manuscript. Our appreciation also goes to all the peer reviewers for their insights and suggestions.

## Ethics approval and consent of participate

In alignment with the Declaration of Helsinki, ethical clearance for this study was granted by the Ethics Committee of the First Affiliated Hospital of Nanjing Medical University (approval number: 2025-SR-718). The animal experiments were approved by the Institutional Animal Care and Use Committee at Yangzhou University (Approval Number: 202607027). All experimental procedures adhered to the “Guidelines for the Care and Use of Laboratory Animals” established by the National Institutes of Health.

## Data Availability Statement

All information produced or examined throughout this investigation, along with the fundamental data underpinning the conclusions of this research, is encompassed within the manuscript. Any additional details or inquiries concerning the data may be addressed to the corresponding author upon a reasonable request.

## Supplemental Material

Expanded Methods

Figures S1-S7

Table S1

